# A novel Boolean network inference strategy to model early hematopoiesis aging

**DOI:** 10.1101/2022.02.08.479548

**Authors:** Léonard Hérault, Mathilde Poplineau, Estelle Duprez, Élisabeth Remy

## Abstract

Hematopoietic stem cell (HSC) aging is a multifactorial event that leads to changes in HSC properties and function. These changes are intrinsically coordinated and affect the early hematopoiesis, involving hematopoietic stem and progenitor cells (HSPCs). The objective of this work is to better understand the mechanisms and factors controlling these changes. We have therefore developed an original strategy to construct a Boolean network of genes explaining the priming and homeostasis of HSCs **(graphical abstract)**. Based on our previous scRNA-seq data, we performed an exhaustive analysis of the transcriptional network and identified active transcription modules or regulons along the differentiation trajectory of selected HSPC states. This global view of transcriptional regulation led us to focus on 15 components, 13 selected TFs (Tal1, Fli1, Gata2, Gata1, Zfpm1, Egr1, Junb, Ikzf1, Myc, Cebpa, Bclaf1, Klf1, Spi1) and 2 complexes regulating the ability of HSC to cycle (CDK4/6 - Cyclin D and CIP/KIP). We then defined the connections controlling the differentiation dynamics of HSC states and constructed an influence graph between the TFs involved in the dynamics by mixing observations from our scRNA-seq data and knowledge from the literature. Then, using answer set programming (ASP) and in silico perturbation analysis, we obtained a Boolean model which is the solution of a Boolean satisfiability problem. Finally, perturbation of the model based on age-related changes revealed important regulations, such as the overactivation of Egr1 and Junb or the loss of Cebpa activation by Gata2, which were found to be relevant for the myeloid bias of aged HSC. Our work shows the efficiency of the combination of manual and systematic methods to elaborate a Boolean model. The developed strategy led to the proposal of new regulatory mechanisms underlying the differentiation bias of aged HSCs, explaining the decreased transcriptional priming of HSCs to all mature cell types except megakaryocytes.

**Graphical abstract:** From single cell RNA-seq (scRNA-seq) data and current knowledge in early hematopoiesis (literature and biological database investigation), 3 inputs were obtained to define the network synthesis as a Boolean Satisfiability Problem depending on observations of states in the differentiation process:

1. Influence graph between selected components.
2. Discretized component activity levels in the considered states (blue: 0/inactive, white: */unknown or free, red: 1/active).
3. Dynamic relations (stable states, (non) reachability) between the considered states. Then, these inputs were encoded as constraints in Answer Set Programing (ASP) thanks to the Bonesis tool. After the solving, a Boolean model of early hematopoiesis is obtained. This model is altered according to the characteristics of aging observed in our scRNA-seq data, in order to identify the main molecular actors and mechanisms of aging.

Graphical abstract:
Overview of the scRNA-seq assisted gene Boolean network synthesis strategy.

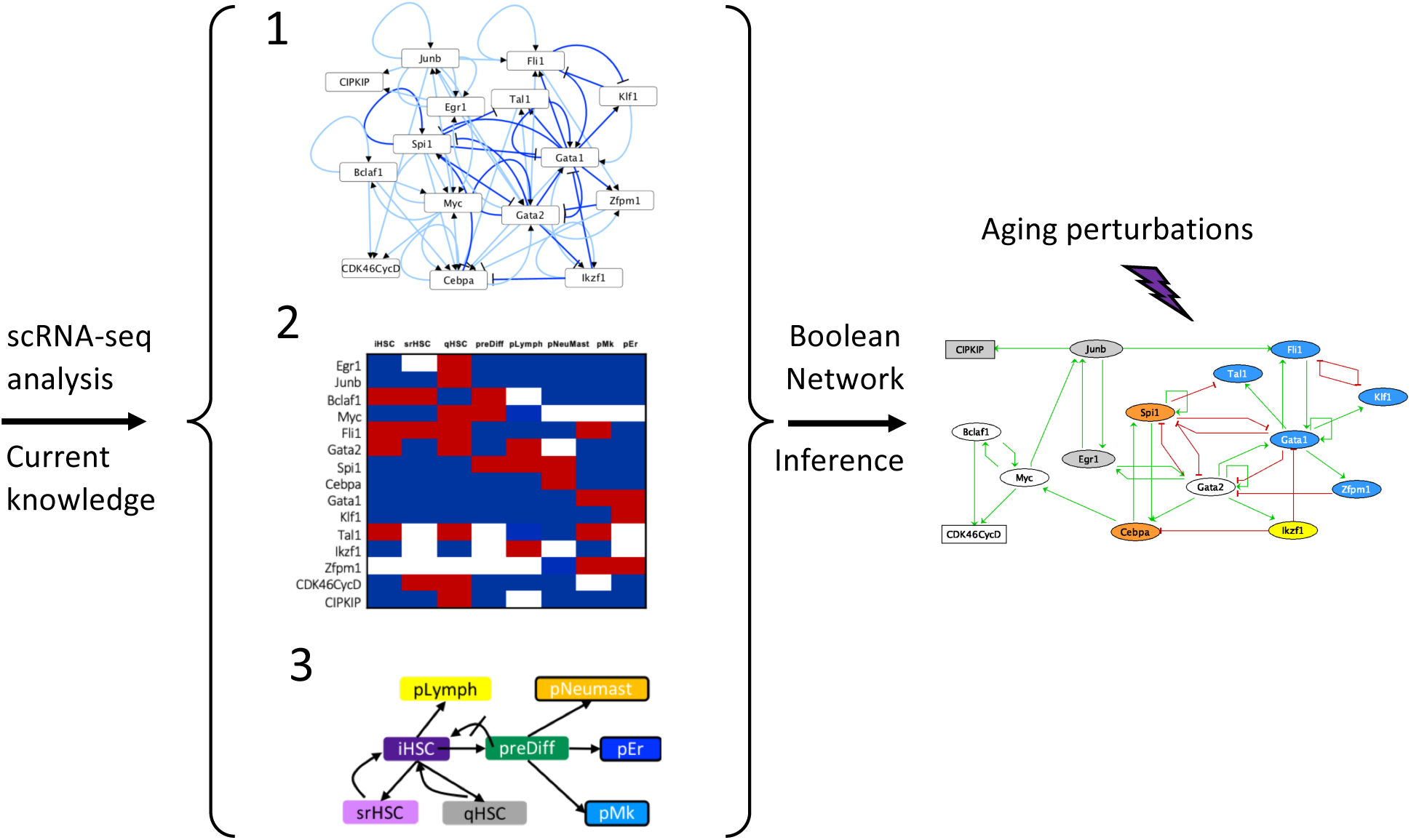

## Introduction

Hematopoiesis is the process of cellular differentiation that allows the hematopoietic stem cell (HSC) to produce all types of mature and functional blood cells. A critical balance between HSC self-renewal and differentiation into different hematopoietic lineages must be maintained throughout an individual’s life in order to maintain an effective immune system and normal oxygen transport. As with many biological systems, transcriptional regulations orchestrated by transcription factors (TFs) and their networking are key mechanisms for instructing differentiation occurring in the hematopoietic stem and progenitor cell (HSPC) compartment (reviewed in ^1,2^). The characterization of their activities has led to refine our view of the multiple branch points for early hematopoiesis^3^.

It is also well known that deregulations of transcriptional and epigenetic programs underlie the decline in HSC function during aging^4,5^. This leads to an alteration of the HSPC pool phenotype, resulting in an increase in myeloid and megakaryocytic cells at the expense of lymphoid and erythroid ones in aged individuals^6,7^. As a consequence, elderly people are subject to various blood disorders such as anemia and acute myeloid leukemia. Given the current aging of the population, deciphering the molecular mechanisms and more particularly the gene regulatory networks (GRNs) underlying age-induced deregulation of HSPCs is of great interest and is the subject of extensive research.

With recent technology developments allowing single-cell resolution transcriptome analysis and lineage tracing, hematopoiesis is now considered as a continuous process with a very early and gradual priming of the HSPC compartment into different lineages^8,9^. Comparative transcriptome studies from young and aged HSPCs at the single-cell level (scRNA-seq) have accurately mapped lineage priming and cell cycle changes in aged mice, allowing the identification of subgroups of HSPCs distinct in their ability to maintain early hematopoiesis and whose proportions are altered during aging^10–12^. With thousands of gene expressions measured in thousands of cells, scRNA-seq also provided the amount of data needed to significantly improve GRN inference methods^13,14^. Some inference methods, based on mutual information or regression trees, have been successfully used to analyze regulatory networks in the HSC microenvironment^15^. They have also permitted the identification of regulons, modules formed by a transcription factor and its target genes, in the HSPC compartment during human ontogeny^16^ or mouse aging^10^. However, these studies only provide a static view of the GRN governing HSC fate. It would be relevant to have a dynamic view of the molecular mechanisms and interactions involved in cellular decisions such as commitment to a particular lineage. To address these issues, one possibility is to study the dynamics of networks using Boolean network (BN) modeling^17^. BN modelling approach provides a good abstraction of the long-term behaviors of a biological system, although continuous changes in component activities and timing of regulations are not captured. It also provides mechanistic explanations on the functioning of regulatory processes without the need for kinetic parameters and is therefore particularly suitable for the analysis of large biological networks^18^. In the context of hematopoiesis, several logical models of HSC differentiation have been proposed, which helped us to understand the connection between the major TFs specifying hematopoietic lineage differentiation^19–21^.

The recent development of scRNA-seq technology has opened up new possibilities and challenges in the field of BN modeling. Indeed, the scRNA-seq data represent observations of a large number of cell states that when ordered along a pseudo-trajectory and after binarization of the component activities of interest (usually gene expression), can be interpreted as an observation of a trajectory generated by a BN. It is then possible from the transitions between states observed in the data to find logical functions for each component by a reverse engineering approach, in view of the observed dynamics^21,22^. More recently a method, called Bonesis, has been developed to infer, from a gene regulatory network, a BN satisfying dynamic constraints between cell states^23^ and it is most likely that scRNA-seq data through the analysis of the pseudo-trajectory are well suited to extract the dynamical constraints used as input for Bonesis.

Based on scRNA-seq analysis, we and others recently observed new priming events in the HSC pool^8,10^ which are deregulated with aging. Here, we wanted to take advantage of our scRNA-seq data of young and aged mouse HSPCs^10^ to construct a BN to understand the dynamic of early priming of HSCs and to precisely characterise transcriptional determinants leading to HSC dysfunctions and aging. We first defined the key states of early hematopoiesis by selecting and grouping HSPCs according to their transcriptome in coherence with their TF marker activity. Next, combining our analyses with the current literature on HSC biology, we adapted an existing GRN of early myelopoiesis^24^, by inferring with Bonesis a BN whose dynamics corresponding to our HSC priming pseudo trajectory. Finally, we performed and analyzed perturbations of this BN to propose key factors and mechanisms supporting the differentiation bias observed in aged HSCs. Overall, our results provide a mathematical model of early hematopoiesis that allows us to assess the regulation of age-related physiological changes in HSC.

## Methods

### scRNA-seq dataset

We used the scRNA-seq dataset presented in our previous study available in the Gene Expression Omnibus database under accession code GSE147729^10^. This dataset is composed of two pools of young (2/3 months) mouse HSPCs and two pools of aged (18 months) mouse HSPCs. Our previous results (cell cycle phase assignment, cell clustering, pseudotime ordering) were considered in this study to define the HSPC states at the basis of our modeling^10^ (**Supplementary Table 1**).

### Regulon analysis

#### Identification of regulons with pySCENIC

We used the Single-Cell Regulatory Network Inference and Clustering (SCENIC) approach^25^ to identify regulons, which are modules of one TF and its potential targets, and their activities. We ran SCENIC workflow using pySCENIC v1.10.0 with its command line implementation^26^ as in our previous study regarding gene filtering, TF motifs (motifs-v9-nr.mgi-m0.001-o0.0), cis-target (+/- 10 kb from TSS mm9-tss-centered-10 kb-7species.mc9nr) databases and command line options^10^. In this work, we used as input all 1721 TFs with a motif available in the motif database. We processed with SCENIC workflow all cells together as well as only young or only aged cells. For each cell set, regression per target step with grnboost2 followed by cis-target motif discovery and target pruning were run 50 times using a different seed for the pySCENIC grn command. The regulons and their targets recovered in at least 80% of the runs were kept. For a gene *g* with n regulators (*r*_1_ …, *r*_*n*_), the normalized interaction score (NIS) of the transcriptional regulation of *g* by *r*_*t*_ is defined as follows:

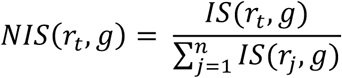

where *IS*(*r*_*t*_.,*g*) is the interaction score defined as the product of the number of SCENIC runs in which the interaction from *r*_*t*_. to *g* was found, by the average importance score given by grnboost2 for the interaction across these scenic runs. The results for the interactions found in pySCENIC analysis of all cells are available in **Supplementary Table 2. Regulon markers of HSPC states**. We scored the activating regulons (i.e., regulons with a positive correlation between the TF and its targets) with AUCell (pySCENIC aucell command, default option with a fixed seed). Averaged AUCell scores by HSPC states were computed. These scores were standardized in order to hierarchically cluster the regulons using ward.D2 method of the R function hclust with Euclidean distance. The DoHeatmap function from the Seurat v3 package^27^, was used to display the results. Averaged AUCell enrichment scores for young and aged cells by HSPC states were also computed in the same way.

Activating regulon markers of HSPC states were identified based on their AUCell scores using FindAllMarkers Seurat function (min.pct=0.1, logfc.threshold=0) with Wilcoxon rank sum tests. Only regulons with an average AUCell score difference above 0.001 between one state versus all the others were kept. A p-adjusted value (Bonferroni correction) threshold of 0.001 was applied to filter out non-significant markers (**Supplementary Table 3**).

Activating regulon activity differences with aging in each state were identified using the FindConservedMarkers Seurat function (sequencing platform as grouping variable, min.pct = 0.1 and logfc.threshold = 0) with Wilcoxon rank sum tests. For each HSPC state, only average AUCell score differences of the same sign and above 0.001 in the two batches presenting a combined p value < 0.001 were kept (**Supplementary Table 4)**.

#### TF network

A network based on interactions between TFs found in at least 90% of SCENIC runs on all cells was built (discarding self-inhibitions because of their uncertainty ^21^). The cluster_louvain function, from igraph R package^28^ was used to find TF communities in the undirected transformation of this network with edges weighted by the NIS scores. The Cytoscape software^29^ was used to visualize the results from graph clustering.

### Cistrome database analysis

Available mouse TF ChIP-seq experiments annotated for bone marrow tissue were analyzed using Cistrome database workflow^30^. More specifically, for each bed file of the selected experiments in the databases, the top 10,000 peaks with more than 5-fold signal to background ratio were conserved for downstream analysis. Then, target transcripts were identified with BETA in each TF experiment^31^. We considered all TF peaks in an experiment *j* inside a +/- 10 kb window from a Transcriptional Start Site (TSS).

BETA gave us a regulatory score *s*_*i*_ for each 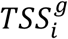 of potential target genes g of *a* TF *t*. Then we defined a global cistrome regulatory score (*CRS*) for a TF *t* on a potential target gene *g* as follows:

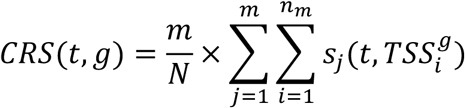

where, the 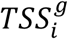 are the *n*_*m*_ TSSs of g for which a regulatory score *s*_*i*_ by *t* is obtained in experiment *j* among the *m* experiments where the regulation is found. This score is weighted by the *m,N* ratio where *N* is the number of experiments for the given TF available in the considered cistrome datasets. Only *CRS* for Scenic interactions or referenced regulations were retained (**Supplementary Tables 2 & 5**).

### Boolean modeling

A **Boolean Network** (BN) is an influence graph parameterized with logical functions. An influence graph is a directed signed graph that is an abstraction of regulatory and molecular interactions (in binary relations). Nodes stand for biological components (here TFs and cell cycle protein complexes) that are connected through edges representing activations and inhibitions. In this study, we constructed an influence graph of 15 nodes and their interactions involved in early hematopoiesis (see results). From this influence graph we define a discrete dynamical model using logical formalism. Each node of the influence graph is associated with a Boolean variable representing its level that can be 0 (component inactive) or 1 (active), this level reflecting its ability to regulate its targets. The effect of regulators on the level of the target node is expressed through **logical functions** (using connectors & for AND, | for OR and ! for NOT). Given a **configuration of the network**, i.e., a vector containing the level of all the components, several components may be called to update their level by the logical functions. The choice of the updating policy defines the trajectories of the system (succession of consecutive configurations). Here, we used the Most Permissive (MP) semantics (that is required to use the Bonesis inference tool). This recently proposed semantic considers additional transient states reflecting increasing (↗) or decreasing (↘) dynamical states. A component in (↗) or (↘) state can be read non-deterministically as either 0 or 1 to take into account the uncertainties of its actual influence thresholds on its different targets. This generates a non-deterministic dynamic with a large set of trajectories, which considerably reduces the complexity of the exploratory analysis of the dynamics^32^.

The **attractors** of the model capture the asymptotic behaviors of the system. They are a set of configurations from which it is not possible to escape, and can be fixed points, i.e., a configuration whose all components are stable, or cyclical attractors, containing more than two configurations among which the system oscillates. Finally, a biological interpretation of these attractors, based on the level of some nodes or read-outs of the model, allows them to be associated with biological phenotypes.

BN offer the possibility to easily simulate **perturbations of gene activity**, such as a gain-of-function or overexpression denoted KI (Knocked In), or a loss-of-function or deletion denoted KO (Knocked Out), by maintaining its variable at 1 or 0 respectively. We can also simulate an edgetic mutation by perturbing not a node but an edge of a network. For that, we removed the edge of the network and updated the logical rules of the target nodes.

### Data discretization

We associated each HSPC state with a vector, called **meta-configuration**, representing the discretized activity level of each of the 15 components of the model. The discretization method depends slightly on the nature of the components.

The TF activities at the head of a regulon with more than 10 targets were discretized with a Kmeans clustering of two on all cell regulon activity scores (the value of the cluster with the most cells was retained). Otherwise, because the AUCell scores were less reliable, we discretized the activity on 3 levels, active (1), inactive (0) or free/unknown (*) with Kmeans clustering (K=3) on averaged RNA levels per HSPC states. This was the case for Tal1, Ikzf1 and Zfpm1. For the cell cycle complexes (CDK4/6CycD and CIP/KIP) we took the discretization of the RNA levels genes coding for their component and attributed them a value in {-1, 0, 1} (−1 meaning inactive, 0 free or unknown, 1 active). If the sum of gene values for each complex was above 1, we considered the complex as active, below -1 as inactive, and between -1 and 1 as a free or unknown complex activity.

In order to be less constraining, we relaxed some component activities by replacing their discretized value 0 or 1 to a free (*) status (according to data and biological assumptions, see results).

### Boolean network inference with Bonesis

The influence graph, meta-configurations and dynamical constraints (expressed for instance in terms of stability or reachability of meta-configurations of the network) were encoded in Answer Set Programming (ASP) language with the Bonesis tool which solves our Boolean satisfiability problem and enumerates all the possible Boolean models that satisfy the constraints in the MP semantic^23^.

To reduce the number of solutions we chose to limit the number of clauses per logical rule to 3, as it is the case for most of the logical rules of gene regulatory BN. The solver Clingo^33^ was used in the inference steps.

A subset of 1000 BNs representing a variety of possible behaviors was selected during the generation of the solution space by the Clingo solver, as reported in^34^. For each of them, in silico KO perturbations were performed one by one in the aim of recovering some mutant phenotypes previously described experimentally. In the same way, in silico KI perturbations for nodes with a TF activity upregulated upon aging were conducted. This provides new mutant constraints matching literature evidence for the following next inference steps (**Supplementary Table 6**).

The influence graph was then pruned by adding two optimizations to reduce the number of possible solutions: in priority a maximization of the confident interaction (see results & **supplementary table 5**) number and then a minimization in the other interaction numbers in the inferred models.

### Dynamical analysis of Boolean networks

Dynamical analysis (e.g. attractors reachability from iHSC state, (un)reachabilities between states) of the inferred Boolean models was done in the MP semantics with the mpbn python package^32^.

### Code availability

All R, python, and ASP codes used in this study are integrated in a global snakemake workflow available at: https://github.com/leonardHerault/scRNA_infer.git.

### Statistics

Statistics were computed with R software v4.0.2. The statistical tests for regulon activity scores were performed with Seurat and are detailed above. In each primed HSPC state and in non-primed clusters gathered, the enrichment of age was tested using a hypergeometric test (phyper R function).

## Results

### Regulon analysis identified distinct HSPC states with specific transcription factor activities and their interactions

In order to get references for establishing dynamical constraints for the inference method of BN, we defined HSPC states. We chose to take into account three layers of information considering that each of them is importantly linked to HSPC functionality: cell cluster identity (a meaningful functional partition of the HSPCs), pseudotime trajectory (a good representation of HSPC priming toward different lineages) and cell cycle phases (distinguishing the dividing HSPCs), retrieved from our previous scRNA-seq analyses^10^. Hence, we defined nine states that were visualized on the pseudotime trajectory (**Figure 1A**). We considered two HSPC states at the beginning of the pseudotime trajectory (pseudotime <2); one was composed of non-cycling cells and was called the initiating HSC state (iHSC) and the other one was composed of cells in the G2/M phase and was then considered as the self-renewing HSC state (srHSC). We considered three states based on their cluster identity; the ifnHSC state gathering cells of the ifn cluster (interferon response signature), a state gathering all cells of the tgf cluster that we named the quiescent HSPC state (qHSC state) as all of the cells (except one) were in G1/G0 phase and the preDiff state gathering the cells of the diff cluster representing and spreading on the core of the trajectory^10^. Finally, we defined four lineage-primed HSPC states based on their position at the terminal branches of the trajectory and their belonging to the lineage-primed clusters: pLymph (primed lymphoid clusters pL1 on branch 2), pNeuMast (primed neutrophils and primed mastocytes clusters gathered together, on branch 4), pEr (primed erythrocytes, on branch 5) and pMk (primed megakaryocytes, on branch 5). The selection of these 9 states, excluding cells from our original data set **(Supplementary Table 1**), resumed the initial (iHSC, srHSC), transient (qHSC, ifnHSC), terminal (pEr, pMk, pNeuMast, pLymph) and branching (preDiff) states of early hematopoieis. Thus, we provide an accurate view of the key states that an HSC can reach during early hematopoiesis.

**Figure 1:**
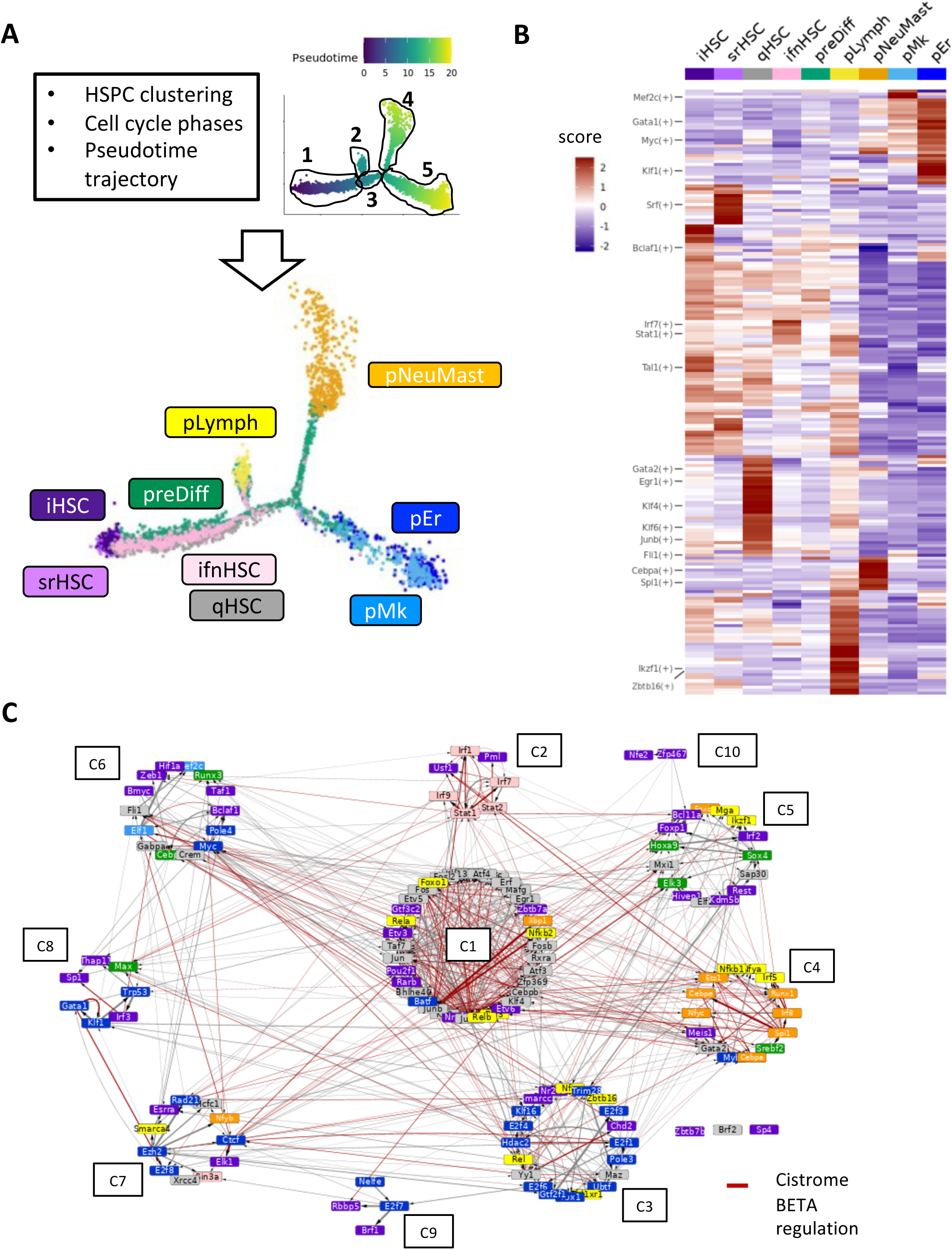
Regulon analysis identified distinct HSPC states with specific transcription factor activities and interactions. **A** Upper panel: HSPC states are defined according to results of cell clustering, cell cycle phase assignment and pseudotime trajectory analysis of scRNA-seq data^10^. On the right, cells are ordered on the pseudotime trajectory and are coloured according to their pseudotime value. The 5 branches of the trajectory are circled. Lower panel: pseudotime trajectory where cells are colored according to their HSPC state: initial HSCs (iHSC, dark violet); self-renewal (scHSC, violet); quiescent (qHSC, gray); interferon (ifnHSC, pink); differentiation (preDiff, green). And the primed states: lymphoid (pLymph, yellow); neutrophils and mastocytes (pNeuMast, orange); erythrocytes (pER, dark blue) and megakaryocytes (pMk, blue). **B** Heatmap of the average AUCell scores of the regulon activity in each HSPC state. The scores were standardized and used to cluster regulons hierarchically. **C** Transcriptional regulation network of the regulon markers of the HSPC states. Regulons were clustered in 10 communities (from C1 to C10) plus 3 isolated nodes with Louvain graph clustering. Node color highlights the states where the regulon is the most active (same color code as in Fig.1 A). Red (resp. grey) edges indicate transcriptional regulations that are (resp. are not) supported by peak analysis in the Cistrome database. Edge thickness represents the normalized interaction score (NIS) obtained from SCENIC.

To functionally characterize these HSPC states, we studied their regulons using the SCENIC workflow^26^. We identified 197 activating and 132 inhibiting regulators **(Supplementary Table 2)**. Among them, 140 were regulon markers of at least one of the 9 HSPC states **(see Methods regulon marker analysis; Supplementary Table 3)**. Next, by quantifying the regulon activities with the AUCell enrichment score^25^ and performing a hierarchical clustering, we revealed a specific regulon activity profile for each of the HSPC states (**Figure 1B**). Regulon activities supported the HSPC state identity. Klf1 was active in pEr, Gata1 in pEr and pMk, Spi1 and Cebpa in pNeuMast and Ikzf1 and Zbtb16 in pLymph. We observed Stat and Irf regulon activity in ifnHSC; Gata2, Junb, Egr1 and Klf1 in qHSC and Bclaf1 and Srf in respectively iHSC and srHSC states. Fli1 regulon was active in both iHSC and pMk. The preDiff state was marked with Spi1 and Myc regulons, two factors involved in HSPC commitment. Except for the srHSC state, uniquely marked by Zbtb7a, probably due to its low cell number, each state was characterized by a combination of regulons, consistent with its transcriptional and lineage feature^10^.

Next, to connect the regulons to each other, we built a transcriptional network whose nodes are TFs at the head of regulons significantly marking at least one of the HSPC states, and directed edges represent the transcriptional regulations between them. When considering only the reliable transcriptional regulations (found in 90% of the SCENIC runs) and after removing auto-regulations, we obtained a directed graph of 133 nodes (TFs) and 670 edges (regulations) (**Figure 1C, Supplementary Table 2**). We further confirmed these regulatory interactions by analyzing the presence of TF peaks in the regulatory regions of their target genes using ChIP-seq data from the Cistrome database^30^. Approximately 60% (302) of the network interactions with an available TF node in the Cistrome database were confirmed by the presence of a peak in the regulatory regions of its targets **(Supplementary Table 2)**. We then performed a clustering analysis by weighting the network using a normalized interaction score (NIS) calculated from the SCENIC results and by applying Louvain clustering. We underlined 10 regulon communities and three isolated regulons (Zbtb7b, Brf2, Sp4). By associating each TF in the network to the most relevant HSPC state, we observed that half of the communities regroups TFs whose activity characterizes the same HSPC state **(Figure 1C and supplementary Table 3)**. Indeed, most of TFs from the C1 community (Klf factors, Jun and Fos AP-1 factors, Egr1) are known to be related to quiescence and their regulons mark the qHSC state, whereas the C2 community contained mainly TFs marking ifnHSC state (eg Irf1-7-9, Stat1-2). In the same way, C3 community was associated with pEr state, C4 community with pNeuMast and C5 with iHSC (Kdm5b, Foxp1) and preDiff (Sox4, Hoxa9) states. It was more difficult to define the smaller communities (C6 to C10) as they presented a more heterogeneous composition of TFs.

Altogether, our analysis revealed a functional relevance of the 9 HSPC states harboring a specific transcriptional activity. We also revealed a well-structured interconnection of TFs that supports the HSC differentiation journey starting with the i-srHSC states, continuing through the transient HSC states (ifn-qHSC, preDiff) and ending with one of the four lineage-primed HSC states.

### Inference of a Boolean network to model HSC priming

To decipher the key molecular mechanisms governing HSC fate, we constructed a Boolean gene network. We developed a strategy based on Bonesis, a recently developed approach for Boolean network inference^23^, which relies on two steps: the synthesis of an influence graph and the definition of dynamical constraints.

**For the influence graph synthesis**, we built a gene network based on a previous published Boolean model of early myeloid differentiation^24^. This model provided megakaryocyte and erythrocyte stable states and a granulo/monocyte branching state that fits well with our defined states^24^. We extracted from it a subgraph of 9 relevant TFs and their mutual interactions. Eight of them were regulon markers of the HSPC states in our analysis: Gata1 a marker of pEr and pMk; Fli1 of pMk; Klf1 of pEr; Spi1 and Cebpa of pNeuMast; Tal1, Fli1 and Gata2 of qHSC (**Supplementary Table 3)**. We also selected Zfpm1, the cofactor of Gata1, which was expressed in pEr and pMk HSPC states (**Supplementary Figure 1**). To adapt the graph to early HSC commitment and aging, we added Ikzf1, a TF involved in early lymphoid specification in HSPC^35^ and whose regulon marked pLymph state according to our analysis (**Supplementary Table 3)**. We also added two components that regulate the ability of HSC to cycle: the CDK4/6-CyclinD (CDK4/6CycD) complex (*Ccnd1-3* and *CDK4/6 genes*) required for the HSC quiescence exit, and its inhibitory complex CIP/KIP (*Cdkn1a-b-c* genes) driving the quiescence of the HSCs^36^. To connect CIP/KIP complex to the transcriptional network, we added Junb and Egr1, two TF involved in HSC quiescence^37–39^, and identified in our regulon analysis as markers of qHSC state and activators of CIP/KIP genes (**Figure 1B; Supplementary Tables 2 & 3**). Finally, to connect CDK4/6CycD to the network, we added Myc and Bclaf1 known to be involved in HSC cell cycle^40,41^. Both were active regulons in the preDiff state whereas only Bclaf1 regulon was active in srHSC state and had CDK4/6CycD complex genes as targets (**Figure 1B; Supplementary Tables 2 & 3**). To connect all these nodes, we considered interactions from the original model^24^ that we complemented with interactions identified by SCENIC and in the literature.

Finally, we obtained an influence graph with 15 components and 60 interactions (**Figure 2Ai**), more than 75% of which were confirmed by at least two of the following information sources; SCENIC, literature or Cistrome (**Supplementary Figure 2A; Supplementary Table 5A, B & C**).

**Figure 2:**
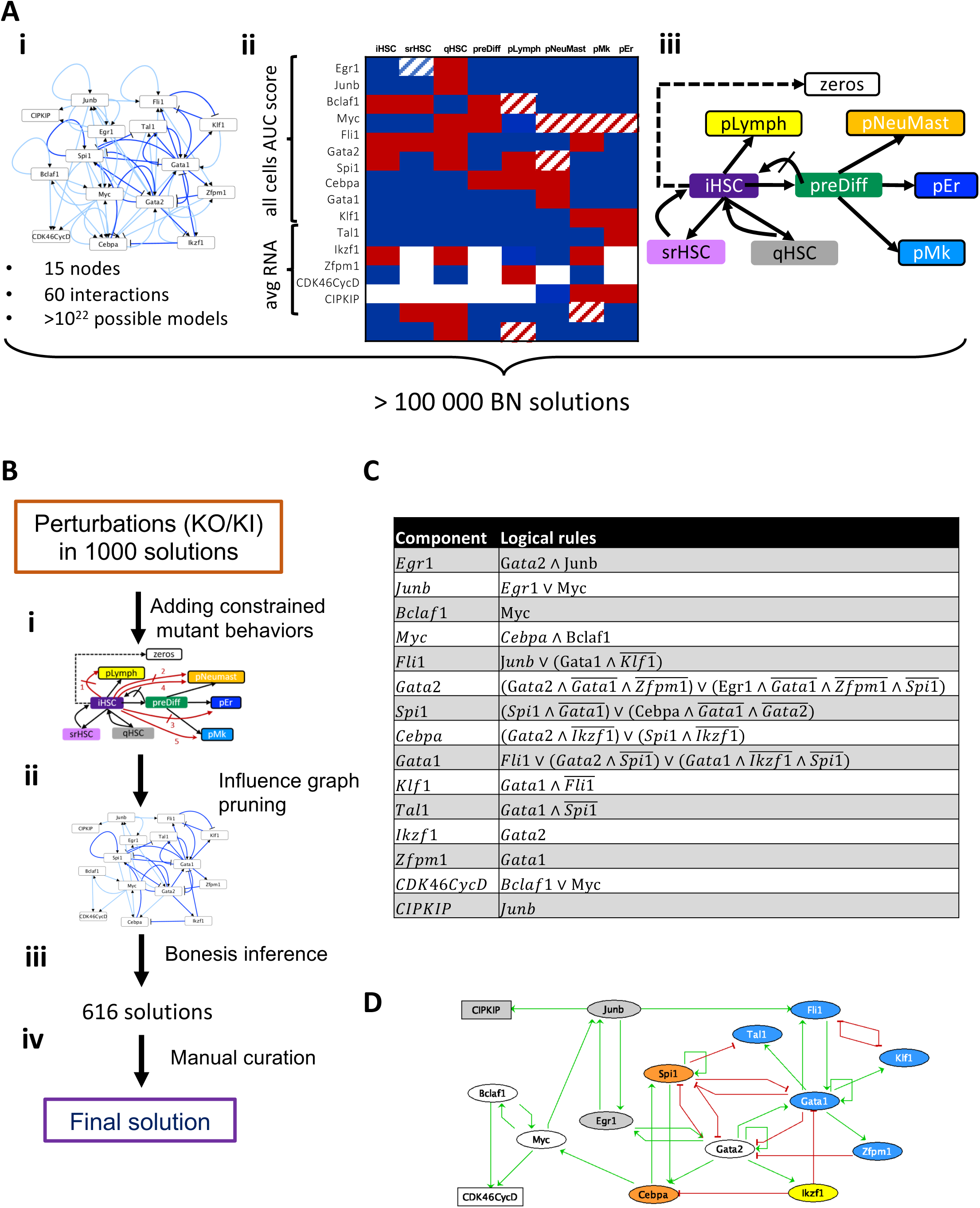
Inference of a gene Boolean network to model HSC priming. **A** Inference steps performed with wild-type constraints. (**i)** A first influence graph is retained taking into account the possible interactions of the components deduced from the literature and the SCENIC results. Interactions with a high (low) confidence level are in dark (pale) blue. (**ii)** Table representing the discretisation of the 15 components in the 9 configurations. Blue indicated active, red inactive and white free state. Red (resp. blue) hatched cases mark node activities freed from 1 (resp 0) to * in the final configuration settings compared to the discretized data. (**iii)** Graph representation of the dynamical constraint imposed between the configurations (nodes). Arrows (resp. crossed out arrows) indicate reachability (resp. unreachability) between source and target configuration. Framed configurations are constrained as fixpoints. Dashed line highlights the reachability of the fixpoint with all node activities at 0 from iHSC. Red (crossed out) arrow highlights the additional (non) reachable constraints of mutant behaviors. **B** Workflow of the strategy used to refine the search of solution and obtain a final solution. **(i)** Updating of the influence graph after the consideration of constraints coming from mutant behaviors. For the updated constraints see **supplementary Figure 3A. (ii)** Pruning of the influence graph through maximization of high-confident interactions and minimization of others. (**iii)** A last inference step enforcing the use of all remaining edges is applied to result in 616 possible solutions. (**iv)** A manual curation is necessary to obtain the final model. **C** Logical rules of the Boolean model. **D** Gene regulatory network of the Boolean model. Nodes, rectangular for cell cycle complexes and ellipse for TFs, are colored according to the HSPC states in which they are highly active according to our regulon analysis: grey for qHSC, yellow for pLymph, orange for pNeuMast, blue for pMk and pEr, white for the nodes highly active in several HSPC states.

For the discretization of the data, we have assigned to each HSPC state a meta-configuration, *i*.*e*., a vector representing the discretized activity level of each of the 15 components (see Methods). This discretization turned out to be too strict regarding the first set of constraints. Thus, we decided to release empirically some constraints on meta-configurations by attributing a free (*) state to some nodes. First, we allowed the cycling configurations CDK4/6CycD in pMk and CIP/KIP in pLymph to be free since they were linked to a HSPC state composed of cells in different cell cycle phases. Following the same idea, we let free the CDK4/6CycD activators, Bclaf1 in pLymph and Myc in pNeuMast, pEr and pMk. We also found that the pNeuMast states presented a bimodal activity for Gata2, with this gene marking pMast and not pNeu cells (see Supplementary Table 1 in ^10^). Thus, we let Gata2 free in pNeuMast HSCP states. Finally, we also let free Egr1 in srHSC in agreement with a previous study suggesting its role in HSC maintenance in the hematopoietic niche^37^.

In order to infer with Bonesis a BN whose dynamics fits with our pseudo-trajectory of HSC priming, we enunciated dynamical constraints between the HSPC states. We required that the model presented at least four fixed points, one for each of the 4 primed meta-configurations pLymph, pNeuMast, pER and pMk, reachable from iHSC. We also added a zeros configuration in which all the components are inactive, reachable only from iHSC directly. We allowed a cell to go back and forth from the iHSC state to the srHSC or qHSC state as suggested by the literature^40,42^. Based on the trajectory and the differentiation committed state of preDiff, we considered this state as a “no return state” and blocked its return to the iHSC state. From this state, any of the 3 primed fixpoints pNeuMast, pER, and pMk were accessible. We allowed a cell from iHSC to directly reach the pLymph fixpoint based on the shape of the pseudo-trajectory and the high hscScore of the pLymph cells^10^. All these constraints are resumed in the HSC differentiation journey **(Figure 2Aiii)**.

With this first inference, Bonesis provided a set of over 100,000 solutions. We therefore developed a strategy to refine the solution search and get a Boolean network solution (**Figure 2B**). We added constraints by considering mutant behaviors: we retained solutions whose mutant simulations agree with the biological phenotypes of these same mutants previously described in the literature (see methods and **Supplementary Table 6** for the references): the Ikzf1 KO conducting to an absence of pLymph fixpoint, the Spi1 KO to absences of both pNeuMast and pLymph fixpoints, the Klf1 KO to an absence of pEr fixpoints and the Junb KO to an apparition of an additional proliferative (active CDK4/6CycD complex) pNeuMast fixpoint. We also considered KI perturbations on Egr1 and Junb nodes, as these components were previously found upregulated in HSC upon aging^11^. For these KIs, we observed a loss of reachability of all fixed points except a quiescent (CIP/KIP active) pMk one consistent with the HSC priming bias we previously described^10^. These 6 altered behaviors are resumed in **Supplementary Figure 3** and were added to the Bonesis constraint set (**Figure 2Bi**). Next, we performed a graph pruning that consists in reducing the number of edges in the influence graph by favoring the most confident ones, which were chosen based on strong literature supports (**Supplementary Table 5A**). There remained 36 interactions (**Figure 2Bii)**, of which more than 80% were supported by at least two sources of information among SCENIC, Cistrome and literature (**Supplementary Figure 2B)**. We required solutions containing all these 36 interactions and another run of Bonesis provided 616 solutions (to compare with the 10^22^ possible solutions with the initial influence graph according to the Dedekind number^43^). These solutions differed on the logical rules of 4 nodes (**Supplementary Table 7**): CDK4/6CycD, Fli1, Gata1 (2 inferred rules for each) and Gata2 (77 inferred rules) and needed a manual curation (**Figure 2Biv**). For the CDK4/6CycD, we chose the rule making the activation possible through Myc in the preDiff state or through Bclaf1 in the srHSC state. For Fli1, we chose the logical rule containing the least number of clauses (2). For Gata1, we chose the rule for which the auto activation is possible only when the two repressors Ikzf1 and Spi1 are inactive. Finally, for Gata2 among the 77 possibilities, 7 contains only two clauses and we chose the one in which the inhibition by Gata1 and its co-factor Zfpm1 are present in both clauses. Finally, we obtained the final Boolean network presented in **(Figure 2C & D)**.

Thus, coupling a customized implementation of Bonesis with multiple sources of biological knowledge allowed to reduce the large number of possible solutions. We successfully synthetized a Boolean network of early hematopoiesis based on a regulatory network consisting of 15 components and 36 interactions.

### Analyses of the Boolean model evidence a sequence of transcriptional events to prime HSCs

Simulations of the model were done within the MP semantic. The dynamics displayed 5 fixed points whose complete descriptions are given **Figure 3A**. According to our requests, all fixed points were reachable from iHSC, regardless of the initial value of the Zfpm1 component. We verified that published mutants related to the genes of the model could be recovered by the simulations. To do this, we simulated the corresponding KO perturbations in the model and compared the results of the simulations to the expected behaviors described in the relevant publications, particularly regarding the reachability of HSPC configurations from iHSC. The large majority of the *in-silico* KO simulations matched the corresponding *in-vivo/in-vitro* perturbations reported in the literature (**Supplementary Table 6**). For example, *in silico Fli1* KO (simulations) led to the loss of pMK fixed point from iHSC in agreement with the *Fli1* KO BM that harbors a megakaryopoiesis defect^44^.

**Figure 3:**
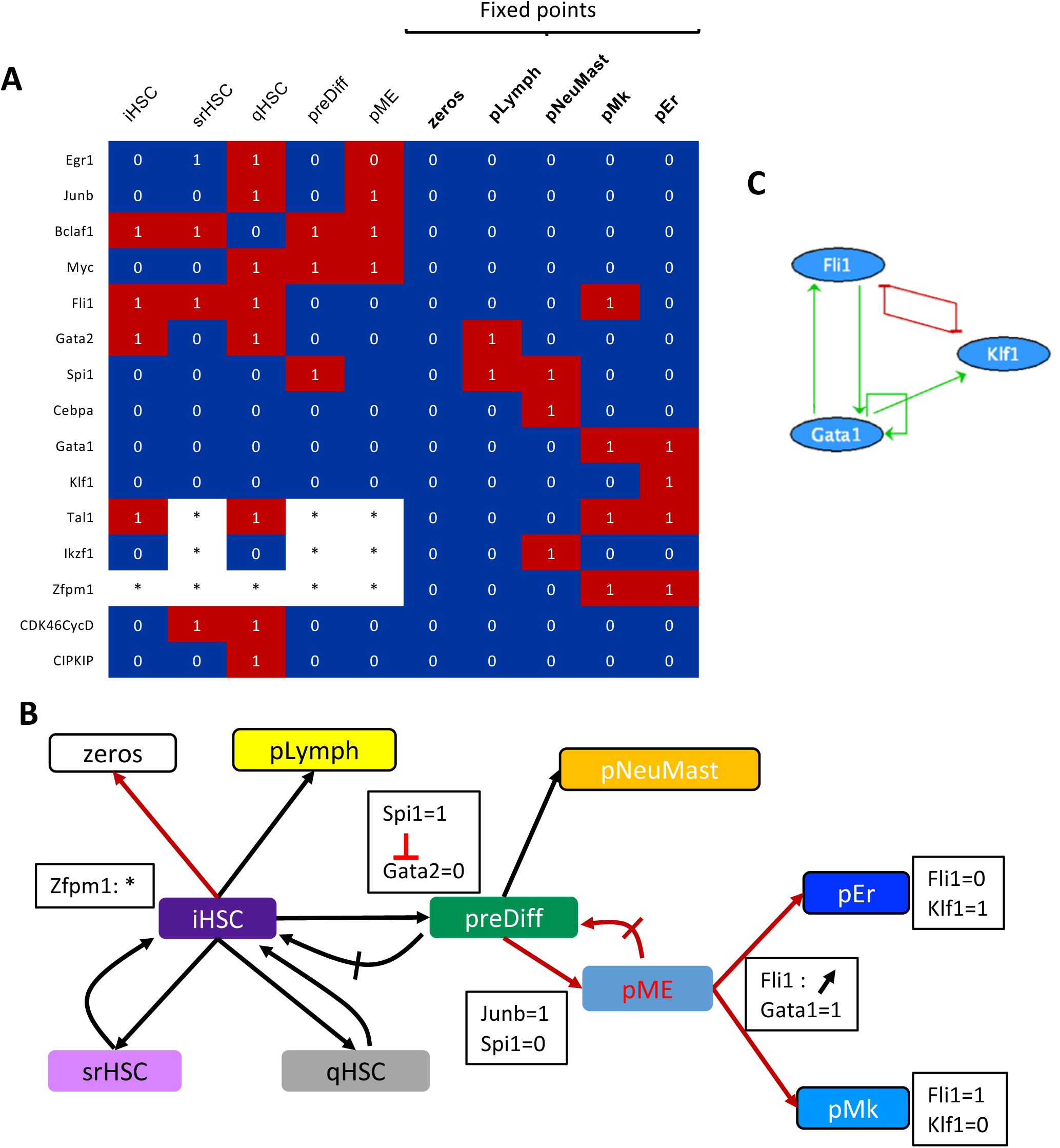
Analyses of the Boolean model evidence a sequence of transcriptional events to prime HSCs. **A** Table describing the states of the model matching the HSPC configurations (column: HSPC states, lines: components of the model). Colors represent the activation levels of the nodes (blue: inactive; red: active, white: free). The five last columns are the stable states of the model. pME configuration (5th column) results from the analysis of the model. **B** Graph representation of the (non)reachabilities between the configurations (nodes). Framed nodes represent fixed points, arrows (resp. crossed arrows) indicate reachability (resp. unreachability) from their source to their target configurations. Black arrows are constrained dynamic properties whereas the red ones result from the dynamic study of the model. Annotations specify some characteristic events (in black boxes): Zfpm1: * highlights the two possible values of this node in iHSC. Irreversible inactivation of Gata2 by Spi1 in the preDiff non-return configuration. necessary update of Junb (=1) and Spi1 (=0) to reach the configuration pME from preDiff. In MP semantics, from pME an increasing activity of Fli1 (↗) can first activate Gata1 and then inhibit Klf1. Thus, depending on whether Gata1 activates Klf1 before it is inhibited by Fli1, pEr is reached rather than pMk. **C** Regulatory motif involving Gata1, Fli1 and Klf1 of the BN with a cross-inhibitory circuit between Klf1 and Fli1 maintaining HSC priming to pMk or pEr.

We conducted a fine analysis of the trajectory space to highlight events that are responsible for some salient dynamical properties along the trajectory. We observed that Gata2 was active in the initial state iHSC (and also in qHSC and pLymph), inactive when the cell reaches preDiff and cannot be re-activated. This event may explain the early branching of the trajectory from the iHSC to the pLymph state, which was characterized by the activity of Ikzf1 whose only regulator is the activator Gata2 (**Figure 3B**). We highlighted a novel transient configuration, named pME and described in **Figure 3A**, that can reach pEr and pMk configurations but not pNeuMast configuration. We observed that the pME configuration was reachable from the preDiff state, when Junb was activated and Spi1 inactivated (**Figure 3B**).

Interestingly, the choice between pMk and pEr fixed points relied on the Fli1-Gata1-Klf1 circuit (**Figure 3C**), and on a transient state of Fli1 caught thanks to the MP semantics. Indeed, starting from the branching point pME in which the three components of the module were absent, Fli1 activity could increase (thanks to the presence of Junb) allowing Gata1 to be also activated and led to the stable configuration pMk. Moreover, as long as Fli1 had not reached its activity level allowing it to inhibit Klf1, activation of Klf1 by Gata1 could occur and led to pEr configuration (**Figure 3B**). It is important to note that we were able to capture this cascade of events thanks to the MP semantics that considers the intermediate states between 0 and 1. It means that a necessary condition to reach pEr from pME is that the inhibition threshold of Klf1 is greater than the activation threshold of Gata1. Finally, the cross-inhibitory circuit involving Fli1 and Klf1 acted as a switch to maintain the differentiation between pMk and pEr (**Figure 3C**).

Furthermore, our model presented a proliferation configuration, in which CDK4/6CycD and Myc were active and CIP/KIP inactive. This configuration was accessible from the iHSC state, and all fixed points could be reached from this state. This is in agreement with our previous results showing an increase in the proliferation of HSC during their priming toward different lineages^10^.

To summarize, the dynamical analysis of our MPBN of early hematopoiesis gives new insights about the succession of early priming events in HSCs. It highlights a decisive role of *Gata2* inactivation to reach preDiff at the expense of pLymph from iHSC. The Spi1 inactivation together with JunB activation are key events to reach from preDiff the pME branching point, whose commitment to the pMK or pER states depends on the fine tuning of Fli1.

### Perturbations of early hematopoiesis model explain some HSC aging features

Our previous single cell RNA-seq analysis revealed an alteration of HSC priming with an accumulation of quiescent HSCs in aged mice at the expense of pLymph, pEr and pNeuMast cells^10^. To decipher the molecular mechanisms and TFs responsible for this alteration, we simulated perturbations in the inferred Boolean network according to alterations observed in the transcriptome of aged HSPCs.

Alterations of regulon activity linked to aging were identified by comparing, for each HSPC state, the regulon activities between young and aged cells. Regulon transcriptional activity differences were found mainly (80%) in the non-primed iHSC, ifnHSC, qHSC states with similar amounts of decrease and increase in activity (**Supplementary Figure 4**), and very few in pNeuMast and pEr. Almost all activity alterations of regulons were found in more than one state (**Supplementary Table 4**). Aging features consisted mainly in a decrease of the activity in regulons related to HSC activation (Runx3, Sox4, Myc and Spi1) and NF-kappaB pathway (Rel and Nfkb factors), and an increase in regulons from the AP-1 complex (Atf, Jun and Fos factors) and involved in quiescence of HSCs (Egr1, Klf factors, Gata2, **Supplementary Figure 5 & Supplementary Table 4**). To be noted that we observed a specific increase in Cebpe-b regulon activity in qHSC state marking the myeloid bias of these quiescent aged cells. Eight of the 13 TF components of our models were altered upon aging in their regulon activities (Myc, Spi1, Junb, Egr1, Fli1, Klf1, Gata2 and Gata1, **Supplementary Table 5B**). More precisely we found Junb, Egr1 and Fli1 (resp. Spi1) activities significantly increased (resp. decreased) in more than a half of the 8 HSPC states considered for the model inference (**Figure 4A**).

**Figure 4:**
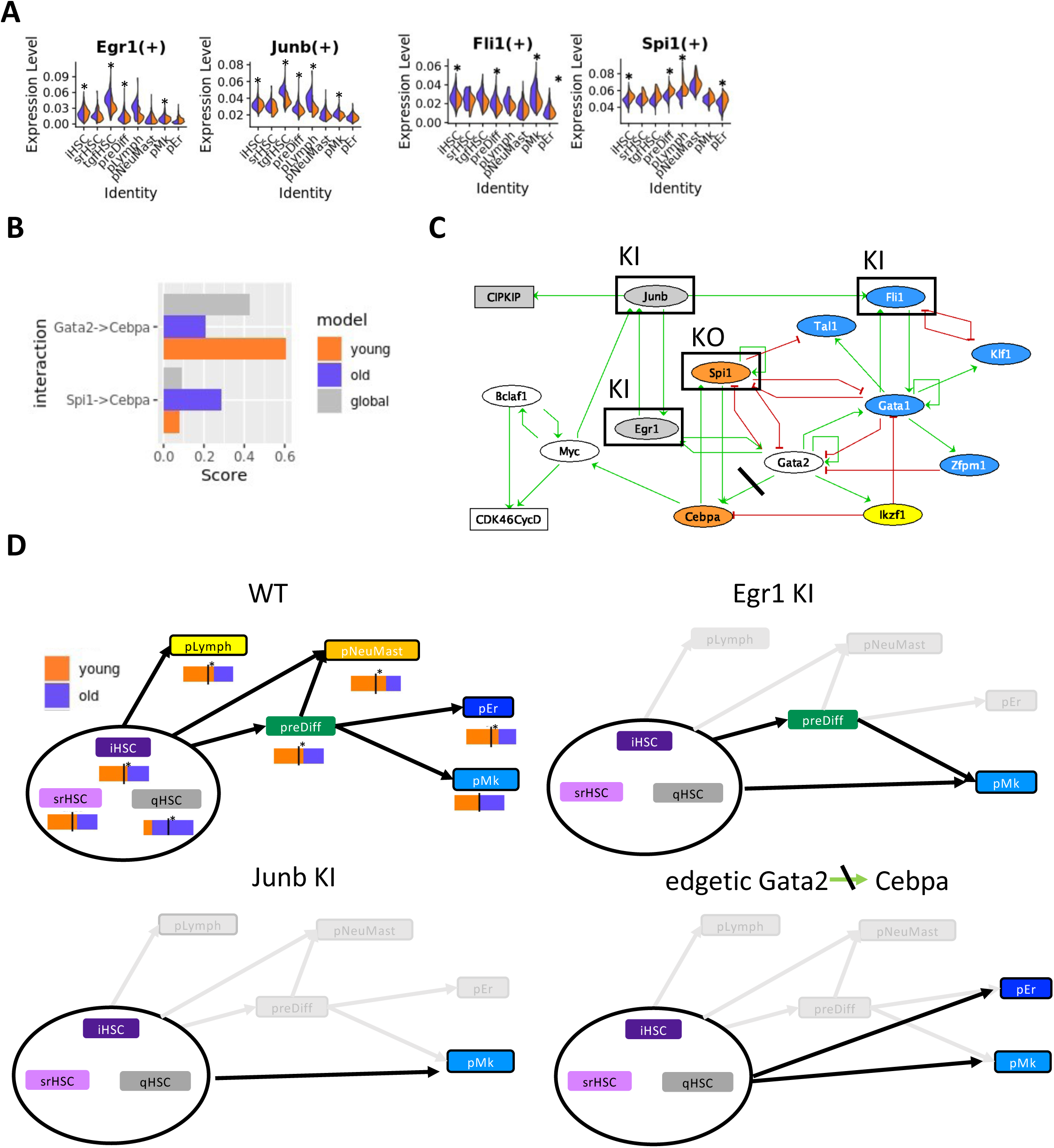
Perturbations of the early hematopoiesis model explain some HSC aging features. **A** Combined violin plots of most altered TF (of the model) activities upon aging in young (orange) and aged (purple) cells from the different HSPC states. Stars show significant differences of activity score between young and aged cells (average difference > 0.001 and p value < 10^−3^). **B** Normalized interaction scores of *Cebpa* activation by Spi1 and Gata2 from SCENIC multiple runs on all cells (grey), young cells (orange) and aged cells (purple). **C** Aging perturbations of the Boolean gene network. Rectangular nodes are cell cycle complexes and ellipse nodes TFs. Nodes are colored according to the HSPC states in which they are highly active according to our single cell analysis: grey for qHSC, yellow for pL, orange for pNeuMast, blue for pMk and pEr, white for the nodes highly active in several HSPC states. Framed nodes highlight the 4 TF significantly altered with aging and crossed out activation of *Cebpa* by Spi1 illustrates its edgetic mutation. **D** Reachability of HSCP states from any initial configuration from iHSC, srHSC or qHSC for WT and 3 altered dynamics of the model: **WT** case (top left) Young (orange) and aged (purple) cell proportion is given below each HSPC state node. A star highlights a significant shift from expected proportion (hypergeometric test p value < 0.05). ***Egr1* KI** perturbation (top right); *Junb* KI perturbation(bottom left); ***Cebpa* edgetic** mutation (bottom right). In each graph, black arrows represent the reachabilities between configurations; pale gray represent the WT reacha bilities lost with the mutation.

To identify possible altered TF regulations, we compared for each regulation the normalized interaction score (NIS) of young and aged cells analyzed separately using SCENIC workflow and computed a score difference between young and aged conditions (**Supplementary Table 4 and Supplementary Figure 6**). The distribution of these score differences showed that most of the regulations were not strongly altered (14% of the interactions have a score difference above 0.4; **Supplementary Figure 7**). When focusing on the interactions of the model supported by SCENIC, we noticed an alteration of *Cebpa* activation by Gata2 (decrease of the NIS of 0.4 upon aging), which was compensated by spi1 activation of C/EBPalpha (NIS increase by 0.2) upon aging (**Figure 4B**).

Thus, in order to simulate aging alteration, we performed KI perturbations on *Junb, Egr1* and *Fli1*, KO perturbation on *Spi1*, and an edgetic mutation (loss of *Cebpa* activation by Gata2) on the network of early hematopoiesis (**Figure 4C**). The simulations of each of these 5 perturbations led to the loss of reachability of the fixed points pLymph and pNeuMast from the i-sr-qHSC configurations. Additionally, the simulation of each of the 3 KIs led to the loss of reachability of pEr, which makes pMK the unique reachable fixed point (**Table 1**). Note that, still for these two perturbations, the constrained fixed point pMk is quiescent (CIP/KIP active). The behaviors of Egr1 KI, Junb KI and Spi1 KO were expected as they were constrained for the inference of the model regarding previous experimental studies (see model inference strategy above). These results agree with our single cell analysis, as the 3 fixed points pLymph, pNeuMast, pEr correspond to the primed HSPC states whose cell proportion significantly decreases with aging, while pMk remains reachable in any of our model perturbations and does not present any decrease in cell proportion with aging in the single cell data (**Figure 4D**). We also observed that preDiff was no longer accessible from i-sr-qHSC configurations with Junb KI disruption and the edgetic Cebpa-Gata2 mutation, suggesting that HSC priming to pMk in aged mice follows an alternative differentiation pathway. This pathway could be directly derived from the state of qHSC cells, the proportion of which increases with aging (**Figure 4D**) and were found at the end of the first branch of the pseudo-trajectory near the appearance of the first pMk (branch 3 and beginning of branch 5 of pseudotime trajectory **Figure 1A**).

**Table 1:**
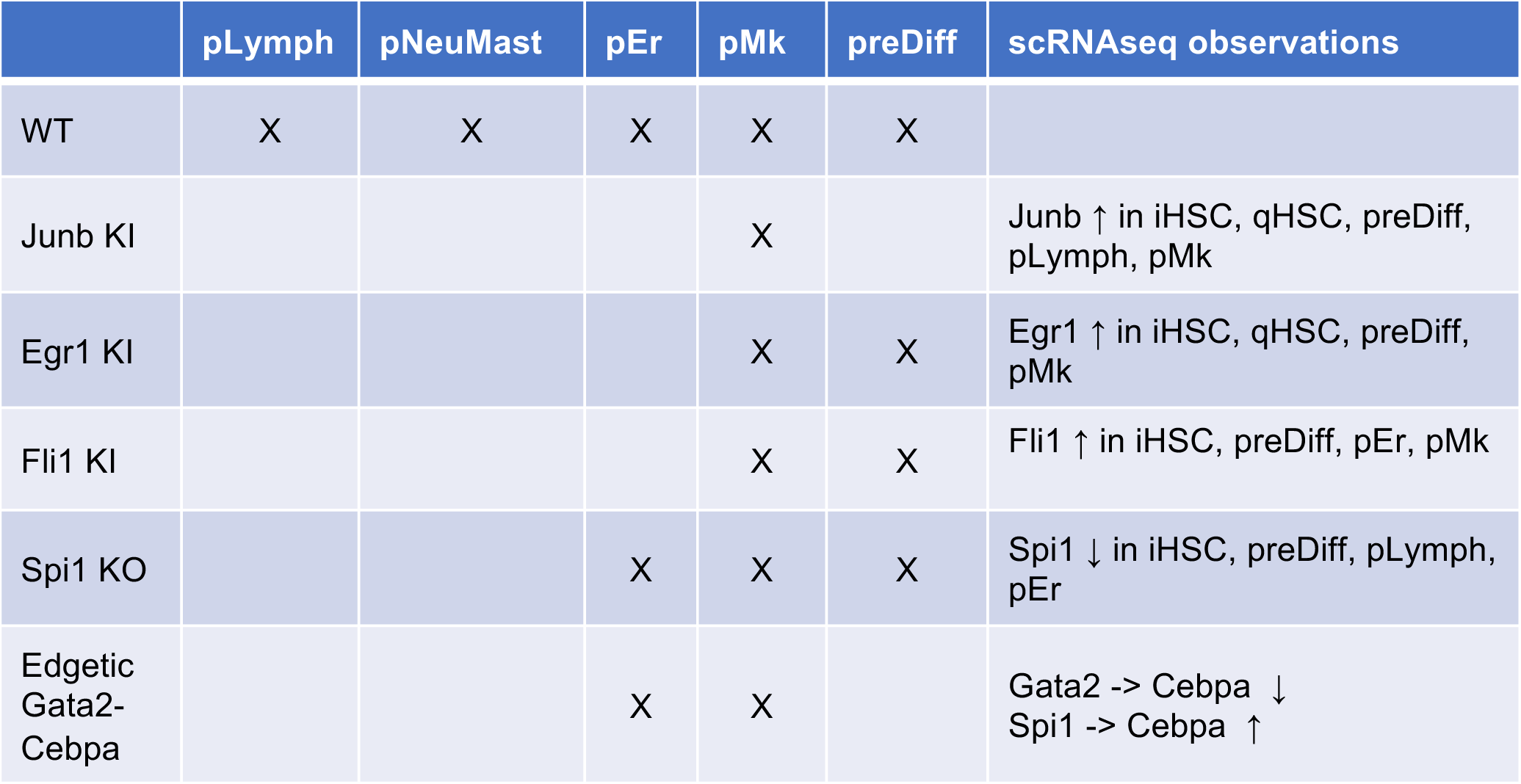
Aging perturbations of the early hematopoiesis Boolean model. This table summarizes the reachabilities of 5 HSPC states (preDiff and 4 fixed points pLymph, pNeuMast, pER, pMk) from states iHSC, srHSC or qHSC, in WT and 5 mutant simulations (a cross indicates that the HSPC-state is reachable). The last column reports the observations in our scRNA-seq data: Up (resp. down) arrows indicate an increase (resp. a decrease) upon aging in component activity or interaction score.

|  | pLymph | pNeuMast | pEr | pMk | preDiff | scRNAseq observations |
| --- | --- | --- | --- | --- | --- | --- |
| WT | X | X | X | X | X |  |
| Junb KI |  |  |  | X |  | Junb ↑ in iHSC, qHSC, preDiff, pLymph, pMk |
| Egr1 KI |  |  |  | X | X | Egr1 ↑ in iHSC, qHSC, preDiff, pMk |
| Fli1 KI |  |  |  | X | X | Fli1 ↑ in iHSC, preDiff, pEr, pMk |
| Spi1 KO |  |  | X | X | X | Spi1 ↓ in iHSC, preDiff, pLymph, pEr |
| Edgetic<br>Gata2-Cebpa |  |  | X | X |  | Gata2 -> Cebpa ↓<br>Spi1 -> Cebpa ↑ |

Therefore, analysis of the model highlighted *Egr1* and *Junb* upregulations and loss of *Cebpa* activation by Gata2 as two major molecular mechanisms that led to HSC aging resulting in the decrease in all lineage priming except the megakaryocyte one.

## Discussion

The mechanisms governing the balance between self-renewal and differentiation of HSCs are the guarantee of functional hematopoiesis, and this balance is altered with aging, Boolean networks provide modelling tools suitable for explanatory analysis of dynamical processes such as the functioning of the HSC differentiation process and the responses to alterations in the interaction network. Several BNs have been proposed to understand the key regulatory elements of HSC differentiation to lymphoid or myeloid progenitors^19–21^ but very few have addressed the impact of aging on HSC properties^45^. Moreover, none of them specifically addressed the early priming of the HSCs, which has been recently emphasized by the development of scRNA-seq analyses^8,10^. While models of BN are classically built from literature, more recent approaches to infer BN from scRNA-seq data have been applied to hematopoiesis^21,22^ albeit presenting some bias due to the imprecision of the pseudotime values due to hidden variables (cell location, epigenetic modification, etc^46^) of scRNA-seq.

In this work, we developed a new modelling strategy that takes full advantage of the scRNA-seq data while incorporating available knowledge from the literature and databases, and that integrates the identified factors playing a role in aging. We obtained a BN explaining the transcriptional mechanisms controlling the priming of HSC toward the different hematopoietic lineages and the impacts of its alterations upon aging. We conducted an original qualitative analysis of the model’s trajectories which consists of tracing succession of events leading to the priming of HSCs in the different lineages. We could propose a main path starting from the initial state iHSC and passing through the intermediate state preDiff that was not captured before. From the iHSC, activation of Ikzf1 by Gata2 stabilizes early lymphoid priming of HSCs, whereas activation of Spi1 and Myc together with an inactivation of Gata2 leads to preDiff state. From preDiff, the positive circuit Gata1/Fli1/Klf1 controls the priming to the neutrophil/mastocytic lineage or the erythroid or megakaryocytic lineages. Our model reproduces the main differentiation trajectory of HSCs under normal conditions, but also shows that alternative trajectories are possible, not passing through preDiff, but allowing direct priming of HSCs towards the different lineages. For example, with early activation of Gata1 by Gata2 the system bypasses preDiff and leads directly to the primed megakaryocyte or erythrocyte state, which is in agreement with previous lineage tracing studies highlighting the coexistence of multiple hematopoietic hierarchies^8,47^. It should be noted that all our analyses were conducted within the MP semantics. This recently defined semantics allows many more transitions between states than the classically used asynchronous or generalized semantics^32^. In particular, our analysis shows that the switch between pEr and pMk from the preDiff state depends on the existence of two thresholds of influence of Fli1 on these targets Klf1 and Gata1, and such a situation cannot be represented with the asynchronous Boolean dynamics. These situations provide perfect case studies to address the question of how the choice of semantics impacts the properties of the dynamics, and more specifically to identify the specific parts of the BN that would need to be refined in order for the MP trajectories to be reproducible in asynchronous semantic.

Our model correctly reproduces behaviors of mutants observed in vivo/in vitro for most of the TFs in the network, except for Myc and Egr1 KO. Simulations of Myc KO showed no difference in silico in the accessibility of primed states. However, an increase in HSC self-renewal and a decrease in differentiation due to intercellular interactions not taken into account in our model have been reported experimentally with this mutation^40^. We also did not observe any changes in dynamics for the Egr1 KO mutant although, again, a previous study showed a decrease in HSC priming along with an increase in HSC self-renewal^37^. These observations could correspond to a transient accumulation of cells prior to delayed priming, not captured by the model.

The loss of lymphoid potential and the myeloid bias, mainly driven by a platelet bias^10,48^ are the main feature of aged HSCs^49–51^. Our model successfully reproduces this aging feature with new molecular mechanisms based on the overactivation of Egr1 and Junb, or loss of Cebpa activation by Gata2. Our results highlight the overactivation of Egr1 and Junb factors in quiescent myeloid HSCs that accumulate with aging^11^, two factors that have been previously involved in HSC quiescence ^37,38^. Our model shows that these alterations impact the positive circuit between Egr1 and Junb required for the multiple HSC priming. Interestingly, the global transcriptional network inferred with SCENIC shows that these two factors are activated by Klf2-4-6 factors known to be downstream of TGF-beta signaling in other biological contexts^52–54^. We saw that aged cells forming the qHSC state have a strong TGF-beta signature combined with a myeloid bias (over-activation of Cebpe-b in particular). Moreover, megakaryocytes are known to promote HSC quiescence by producing TGF-beta^55,56^. Our results therefore sustain the hypothesis of a self-activating loop of HSC aging which would be triggered by TGF-beta increase in aged HSC microenvironment favoring a myeloid-biased quiescent state from which a single priming towards the megakaryocytic lineage would occasionally be possible^57^. Our model proposed a path sustaining this single priming (without passing through preDiff), in agreement with a direct HSC differentiation into megakaryocytes^8^ that would therefore be the one that is preserved by aged quiescent HSCs. Besides, our model shows that the loss of Cebpa activation by Gata2 may drive the loss of lymphoid and neutrophil/mastocyte priming. Knowing the impact of aging on epigenetic^58^, this edgetic alteration could have an epigenetic origin, due to changes either in histone marks on the regulatory elements of Cebpa^59^, or in hypermethylation of the Cebpa promoter as found in leukemia cases^60^.

Thus, we propose a novel model of the intracellular transcriptional network explaining the HSC early differentiation and its megakaryocytes bias related to aging. Analysis of this model could be quantitatively refined to reproduce the probabilities of the different HSC priming observed in the single cell data^61^. More globally, this model could be the basis of a modelling at the cell population level taking into account the HSC and its microenvironment, which is known to be important for HSC aging.

## ACKNOWLEDGMENTS

The authors would like to thank S. Chevalier and L. Paulevé for their assistance with Bonesis.

## AUTHOR CONTRIBUTIONS

E.D. and E.R. conceived and supervised the study. L.H. developed and designed the analyses. L.H. and E.R. analysed and interpreted the data. M.P. and E.D. brought biological expertise. L.H., E.D. and E.R. designed the figures and wrote the manuscript. All co-authors proofread the manuscript.

## Supplementary figure legends

**Supplementary Figure 1: Discretization of gene expressions for the two complexes and regulons with less than 10 targets**. Results of K-means clustering on averaged RNA levels of the selected HSPC states. Blue: inactivated; white: unknown/free; red: activated.

**Supplementary Figure 2: Venn diagrams of influence graph interaction sources. A** The initial influence graph interactions retrieved from SCENIC results and/or literature investigation and supported or not by the Cistrome database analysis (see Supplementary Table 3). **B** The influence graph after pruning.

**Supplementary Figure 3: Constraints and discretization used for the influence graph pruning and the final rule inference. A** Dynamical constraint used for the update of the influence graph and the final rule inference. Black arrows (resp. crossed out arrows) indicate reachability (resp. unreachability) between source and target configuration. Framed configurations are constrained as fixpoints. Dashed line highlights the allowed reachability of a fixpoint with all node activities at 0 from iHSC. Red (crossed out) arrow highlight the additional (non)reachable constraints of mutant behaviors: loss of pLymph reachability with *Ikzf1* or *Spi1* KO, loss of pNeuMast reachability with *Spi1* KO; loss of pEr reachability with *Klf1* KO; additional pNeuMast cycling fixpoint (G2MpNeuMast) with *Junb* KO; a unique pMk quiescent fixpoint with *Junb*/*Egr1* KI (G0pMk). **B** Discretization of component activities in the configurations used for the pruning of the influence graph and the final rule inference. Blue: inactivated (0); white: free (*); red: activated (1). G0pMk and G2MpNeuMast configurations were defined following the first solution space exploration.

**Supplementary Figure 4: Distribution of differences in regulon activities during aging in the different HSPCs states**. Up (Down) marks significant increases (decreases) in activity upon aging (average differences > 0.001, p-value < 10^−3^). An alteration of a regulon activity can be recovered in several HSPC states.

**Supplementary Figure 5: Heatmap of AUCell scores of regulon activities averaged by group of cells from the HSPC states in young and aged cells**. Scores are standardized on aged (A) and young (Y) cells of the different states. Rows are ordered as in Figure 1.

**Supplementary Figure 6: Repartition of the SCENIC interactions retrieved from the analysis of all cells in aged or/and young cells only analysis**.

**Supplementary Figure 7: Histogram of normalized interaction score (NIS) differences with aging**. Interactions found only in all cells analysis with SCENIC have a null difference.

## Supplementary table legends

**Supplementary Table 1: Definition of the HSCP states**. Nine HPSC states were defined according to the results of cell clustering, cell cycle phase assignment and pseudo-trajectory analysis ^1^. Cell number and cell proportion given the entire scRNA-seq dataset are given for each state.

**Supplementary Table 2: Transcriptional network inferred with SCENIC**. The table gives all the transcriptional interactions recovered in at least 80% (40) runs of SCENIC on all cell dataset from a TF head of a regulon toward a target gene with a mor (mode of regulation) of 1 for activation and -1 for inhibition. The recoveredTimes columns give the number of SCENIC runs in which the regulation is recovered. NIS: Normalized Interaction Score computed from importance score of SCENIC. NIS_diff: Normalized Interaction Score differences between NIS obtained from aged cell analysis versus NIS obtained from young cell analysis. NIS difference is 1 when the interaction is recovered in young (resp. aged) cell analysis and not in aged (resp. young) cell analysis. NIS is 0 when the interaction is recovered neither in young or aged cell analysis and only in all dataset analysis. Cistrome_BM column indicates if some ChIP-seq experiments in the Cistrome database for the TF head of regulon were available and analyzed (TRUE) or not (FALSE) enabling the computation of the Cistrome Regulatory Score (CRS) for the interaction.

**Supplementary Table 3: Regulon activity markers of HSPC states**. For each HSPC state, list of the regulons with an average AUCell score difference (avg_diff one state vs all others) > 0.001, a p value (p_val Wilcoxon rank sum test) and a p-adjusted value (p_val_adj Bonferonni correction) < 0.05. Regulons were assigned to a community (C1 to C10) of the TF network.

**Supplementary Table 4: Regulon markers of aging in the different HSPC states**. In each HSPC state Wilcoxon Rank sum tests were performed on the AUCell activity scores between young versus aged cells in batch A and B separately. Only regulons with an activity in at least 10% of either young or aged cells of the state in both batches were tested. The two p-values for each regulon were combined using the Tipett’s method (minimum_p_pval column). In each cluster, the only regulon differences presenting the same variation in the two batches, with an average score difference > 0.001 and a combined p value < 0.001 were kept.

**Supplementary Table 5: Influence graph interactions. A** Interactions between the 15 components considered in the influence graph. List of the transcription factors (TFs) with their mode of regulation (mor: 1 for activation, -1 for inhibition) of a target. When available, references characterizing experimentally the interaction are given. In that case the interaction proof level can be a transcriptional regulation, a physical protein-protein interaction (physical interaction), or a functional interaction: Knock Down (KD), KO (Know Out), retrieved in the specified cell line (cell_line) and or cell type/tissue (cell_type_tissue). The interactions were reliably identified by SCENIC (present in more than 90% of the runs) analysis or not and when it was possible a Cistrome Regulatory Score (CRS) was computed. For cell cycle complexes (CIP/KIP, CDK4/6-CycD) the CRS is the sum of the CRS of each regulation of a considered TF toward one of the genes of the complex. A confidence level of A (high) or B (low) was given depending of references information and CRS and NIS (see **B**) score. After the pruning of low confidence level interactions 36 interactions remained in the solution (solution = TRUE). **B** SCENIC interactions considered for the influence graph. The table gives all the transcriptional interaction recovered in at least 90% (45) runs of SCENIC on all cell dataset from a TF head of a regulon toward a target gene with a mor (mode of regulation) of 1 for activation and -1 for inhibition. The recoveredTimes(_young/_aged) columns give the number of SCENIC runs on all dataset (young dataset/aged dataset) in which the regulation is recovered. NIS(_young/_aged): Normalized Interaction Score computed from importance score of SCENIC on all dataset (young dataset/aged dataset). NIS_diff = NIS_aged – NIS_young. NIS_diff is set to 0 when the interaction is recovered neither in young or aged cell analysis and only in all dataset analysis. Cistrome_BM column indicates if some ChIP-seq experiments in the Cistrome database for the TF head of regulon were available and analyzed (TRUE) or not (FALSE) enabling the computation of the CRS: Cistrome Regulatory Score for the interaction. After the pruning of low confidence level interactions 36 interactions remained in the solution (solution = TRUE). **C** List of all the references used to support some interactions.

**Supplementary Table 6: Comparison of *in silico* KO perturbations in the final BN selected with previous *in vivo/in vitro* mutant studies**. For each altered gene (KO perturbation), the non-reachability of fixed points from iHSC in the perturbed BN was assessed. References presenting in vivo/in vitro related experiments are provided. Some additional fixed points compared to wild type conditions were found (column Remarks). The last column precises if the perturbed behavior is constrained in the inference process (only the mutant behaviors observed in the 1000 selected BN solutions are constrained for the inference of the final solution).

**Supplementary Table 7: Possible rules for CDK46CycD, Fli1, Gata1 and Gata2 after the final inference**. The rules manually selected to select a final solution are in red. Of the 77 possible rules for Gata2 only the 7 ones with two clauses are presented.

