## Supplementary figures 1 to 7 for "A novel Boolean network inference strategy to model early hematopoiesis aging"

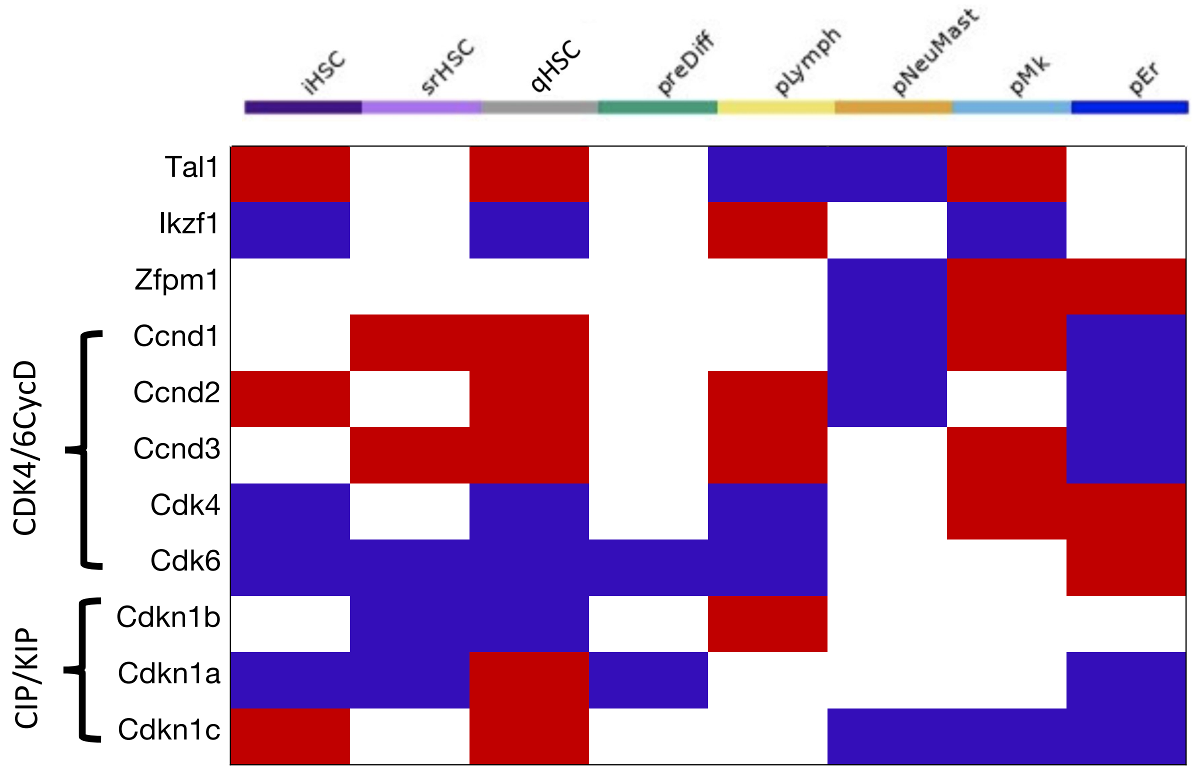


**Supplementary Figure 1: Discretization of gene expressions for the two cycle-related complexes and regulons with less than 10 targets.** Results of K-means clustering on averaged RNA levels of the selected HSPC states. Blue: inactivated; white: unknown/free; red: activated.

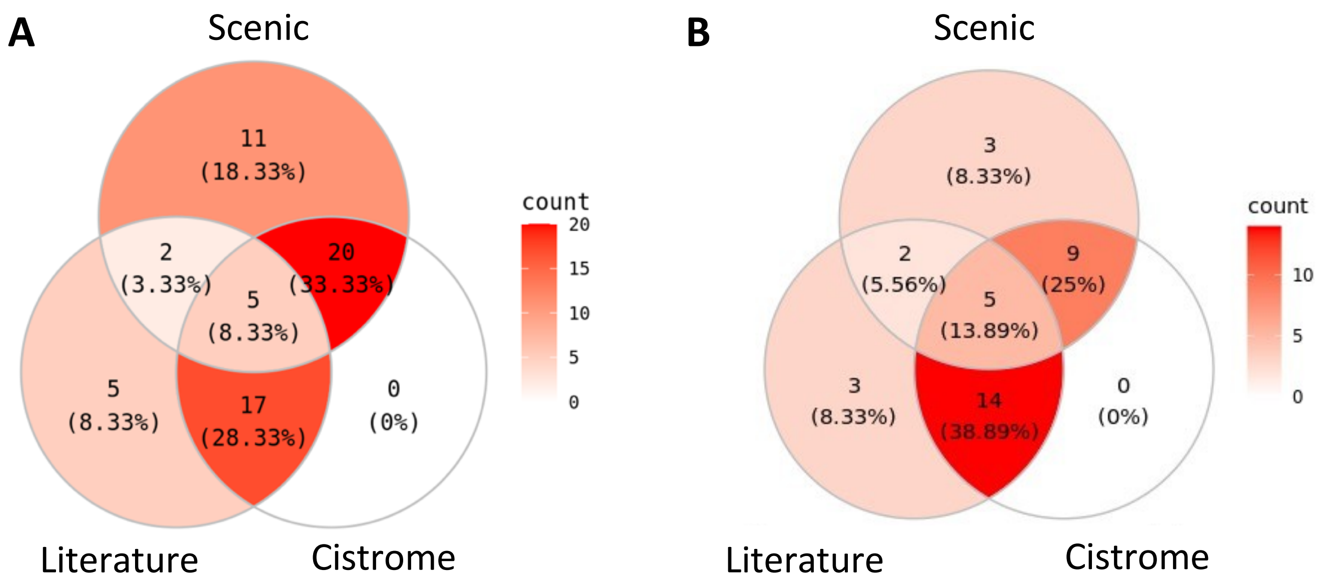


**Supplementary Figure 2: Venn diagrams of influence graph interaction sources. A** The initial influence graph interactions retrieved from SCENIC results and/or literature investigation and supported or not by the Cistrome database analysis (see Supplementary Table 3). **B** The influence graph after pruning.

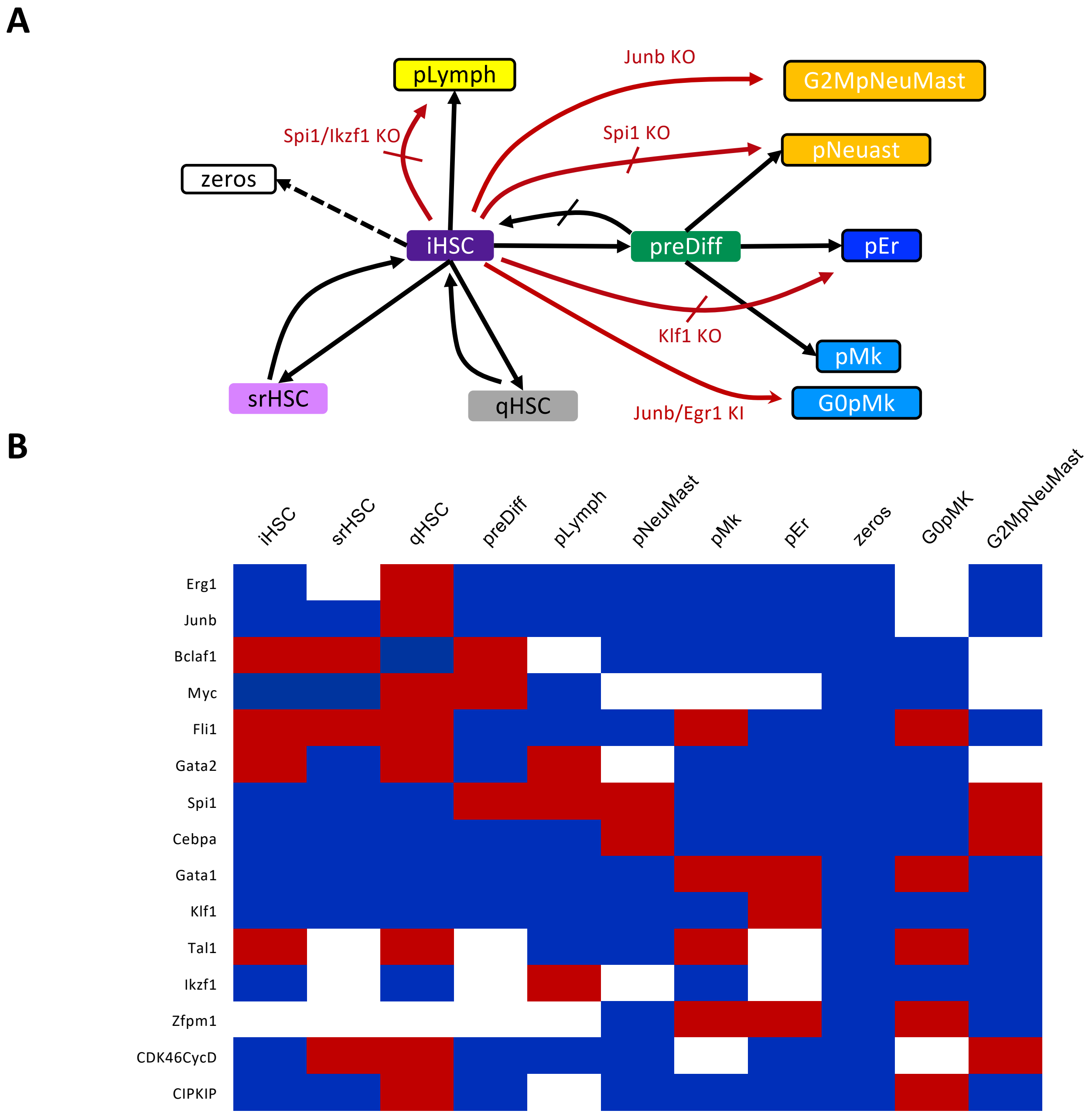


**Supplementary Figure 3: Constraints and discretization used for the influence graph pruning and the final rule inference. A** Dynamical constraint used for the update of the influence graph and the final rule inference. Black arrows (resp. crossed out arrows) indicate reachability (resp. unreachability) between source and target configuration. Framed configurations are constrained as fixpoints. Dashed line highlights the allowed reachability of a fixpoint with all node activities at 0 from iHSC**.** Red (crossed out) arrow highlight the additional (non)reachable constraints of mutant behaviors: loss of pLymph reachability with *Ikzf1* or *Spi1* KO, loss of pNeuMast reachability with *Spi1* KO; loss of pEr reachability with *Klf1* KO; additional pNeuMast cycling fixpoint (G2MpNeuMast) with *Junb* KO;a unique pMk quiescent fixpoint with *Junb/Egr1* KI (G0pMk). **B** Discretization of component activities in the configurations used for the pruning of the influence graph and the final rule inference. Blue: inactivated (0); white: free (*); red:activated (1). G0pMk and G2MpNeuMast configurations were defined according to the first solution space exploration.

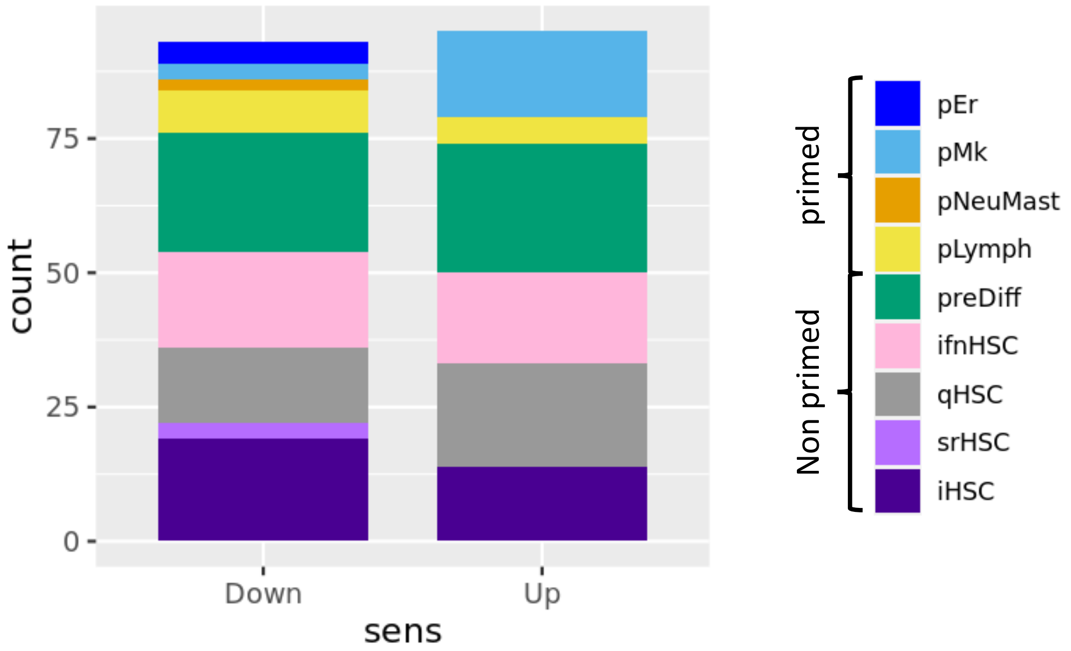


**Supplementary Figure 4: Quantification of regulon activity changes upon aging per state.** Up (Down) marks significant increases (decreases) in activity upon aging (average differences > 0.001, p-value < $10^{-3}$). An alteration of a regulon activity can be recovered in several HSPC states. States are indicated on the right

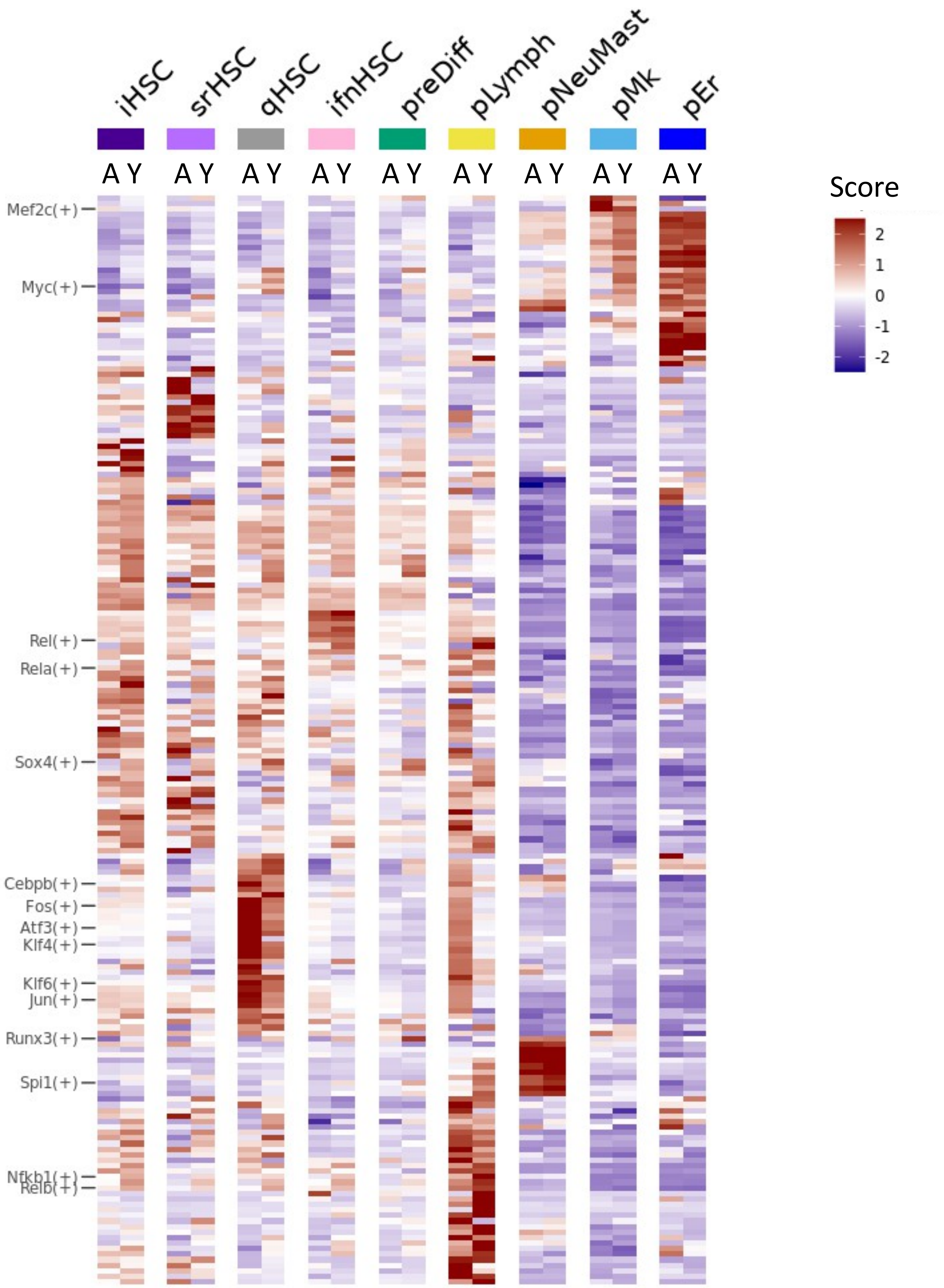


**Supplementary Figure 5: Heatmap of AUCell scores of regulon activity averaged by group of cells from the HSPC states in young and aged cells.** Scores are standardized on aged (A) and young (Y) cells of the different states. Rows are ordered as in Figure 1.

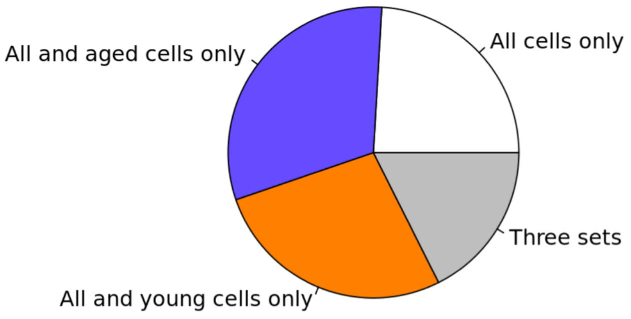


**Supplementary Figure 6:** Repartition of the SCENIC interactions retrieved from the analysis of all cells in aged or/and young cells only analysis.

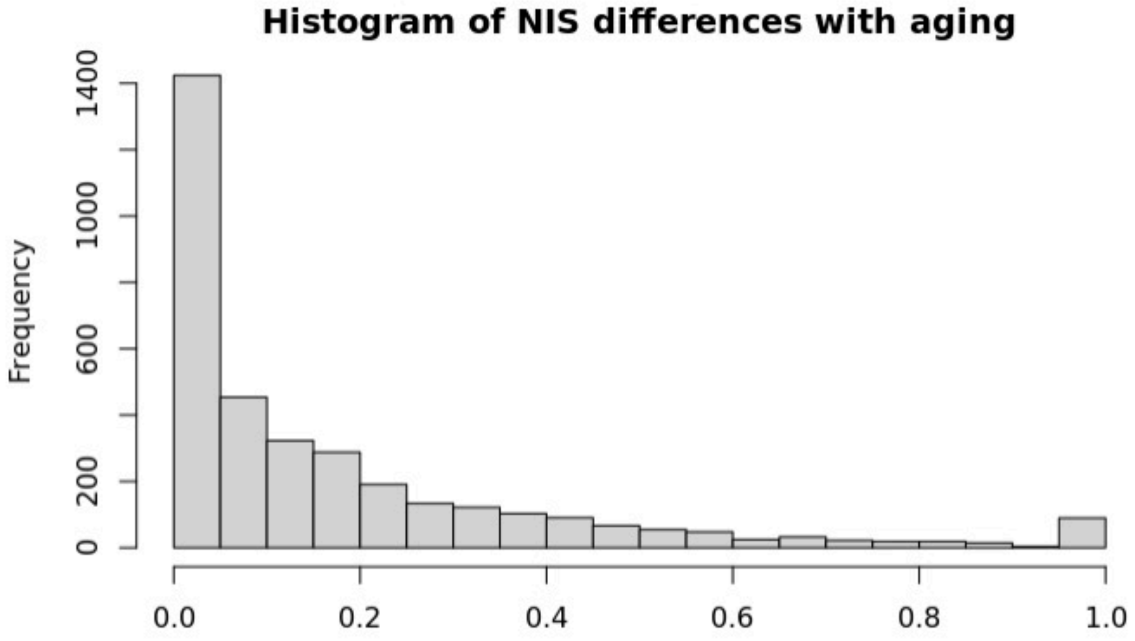


**Supplementary Figure 7 : Histogram of normalized interaction score (NIS) differences with aging.** Interactions found only in all cells analysis with SCENIC have a null difference.
